# In silico investigation of the new UK (B.1.1.7) and South African (501Y.V2) SARS-CoV-2 variants with a focus at the ACE2-Spike RBD interface

**DOI:** 10.1101/2021.01.24.427939

**Authors:** Bruno O. Villoutreix, Vincent Calvez, Anne-Genevieve Marcelin, Abdel-Majid Khatib

## Abstract

SARS-CoV-2 exploits angiotensin-converting enzyme 2 (ACE2) as a receptor to invade cells. It has been reported that the UK and South African strains may have higher transmission capabilities, eventually due to amino acid substitutions on the SARS-CoV-2 Spike protein. The pathogenicity seems modified but is still under investigation. Here we used the experimental structure of the Spike RBD domain co-crystallized with part of the ACE2 receptor and several in silico methods to analyze the possible impacts of three amino acid replacements (Spike K417N, E484K, N501Y) with regard to ACE2 binding. We found that the N501Y replacement in this region of the interface (present in both UK and South African strains) should be favorable for the interaction with ACE2 while the K417N and E484K substitutions (South African) would seem unfavorable. It is unclear if the N501Y substitution in the South African strain could counterbalance the predicted less favorable (regarding binding) K417N and E484K Spike replacements. Our finding suggests that, if indeed the South African strain has a high transmission level, this could be due to the N501Y replacement and/or to substitutions in regions outside the direct Spike-ACE2 interface.

**Hihglights:**

- Transmission of the UK and possibly South African SARS-CoV-2 strains appears substantially increased compared to other variants
- This could be due, in part, to increased affinity between the variant Spike proteins and ACE2
- We investigated in silico the 3D structure of the Spike-ACE2 complex with a focus on Spike K417N, E484K and N501Y
- The N501Y substitution is predicted to increase the affinity toward ACE2 (UK strain) with subsequent enhanced transmissibility and possibly pathogenicity
- Additional substitutions at positions 417 and 484 (South African strain) may pertub the interaction with ACE2 raising questions about transmissibility and pathogenicity

## 1. Introduction

Since the occurrence of the coronavirus disease 2019 (COVID-19), a virulent disease mediated by SARS-CoV-2 initially identified in China, COVID-19 has provoked 2,084,480 deaths with 97,350,867 cases as visualized on Jan 21, 2021 (daily online worldwide data about Covid-19: https://www.worldometers.info/coronavirus/). Although the pathogenesis of this disease remains unclear, in addition to the host response mediated by SARS-CoV-2, variations in the viral strains seem to be also involved in differences in transmission/infectivity and/or severity of the disease. SARS-CoV-2 is a large, enveloped, single-stranded positive-strand RNA virus containing four major structural proteins namely Spike (S) protein, Nucleocapsid (N) protein, Envelope (E) protein, and Membrane (M) protein. The N protein with multifunction is involved in the virus replication, transcription, and assembly and physically interacts with the viral membrane protein during virion assembly [1]. The S protein is a large oligomeric transmembrane protein that mediates the virus entry into host cells. The S protein is composed of two subunits, namely S1 responsible for receptor binding and S2 that mediates downstream membrane fusion **(Figure-1)** [2,3].

**Figure 1.**
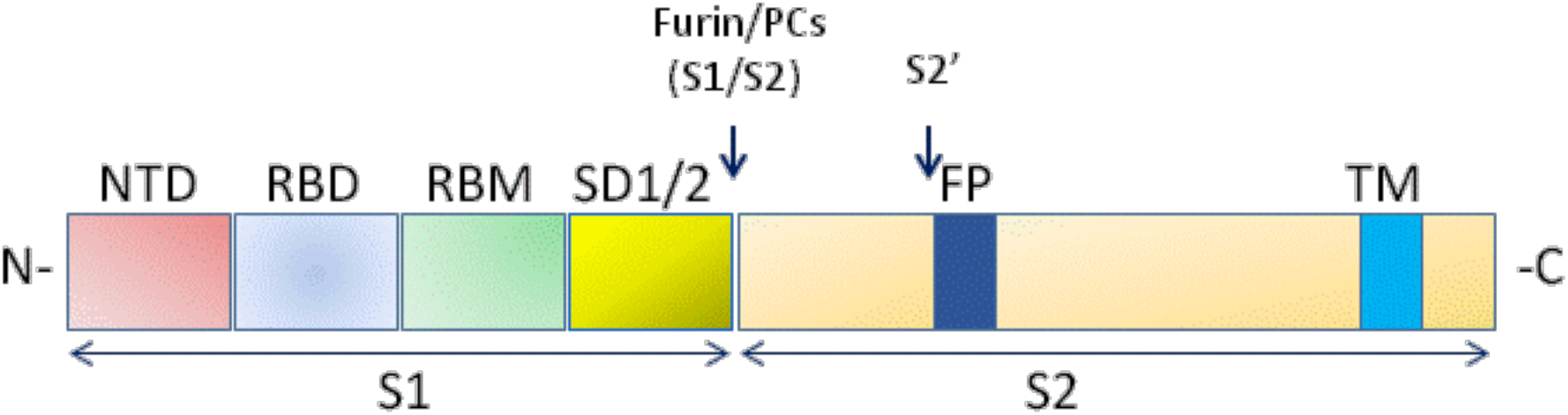
Schematic domain representation of spike glycoprotein, including functional domains in S1 subunit (NTD, N-terminal domain; RBD, receptor-binding domain; RBM, receptor-binding; SD1/2: subdomain 1 and 2) and in S2 subunit (FP, fusion peptide; TM, transmembrane domain. The N and CT terminal domains are indicated. Arrows denote the protease cleavage sites. PCs: Proprotein convertases.

For entry into target cells, SARS-CoV-2 exploits the angiotensin-converting enzyme 2 (ACE2) as a receptor **(Figure 2)** [4]. Therefore, the S protein determines the infectivity of the virus and its transmissibility in the host [5]. Indeed, a small, isolated folded domain of the S1 subunit was reported as the receptor-binding domain (RBD), directly interacts and binds ACE2 during the virus contact with the target cell [6]. This interaction occurs following S protein cleavage at the S1–S2 junction by a furin-like proprotein convertase (PCs) expressed by the host cells [6]. The S proteins of both viruses are further processed in the target cell within the S2 domain at the S2ʹ site, a process that is also necessary for efficient infection [7].

**Figure 2.**
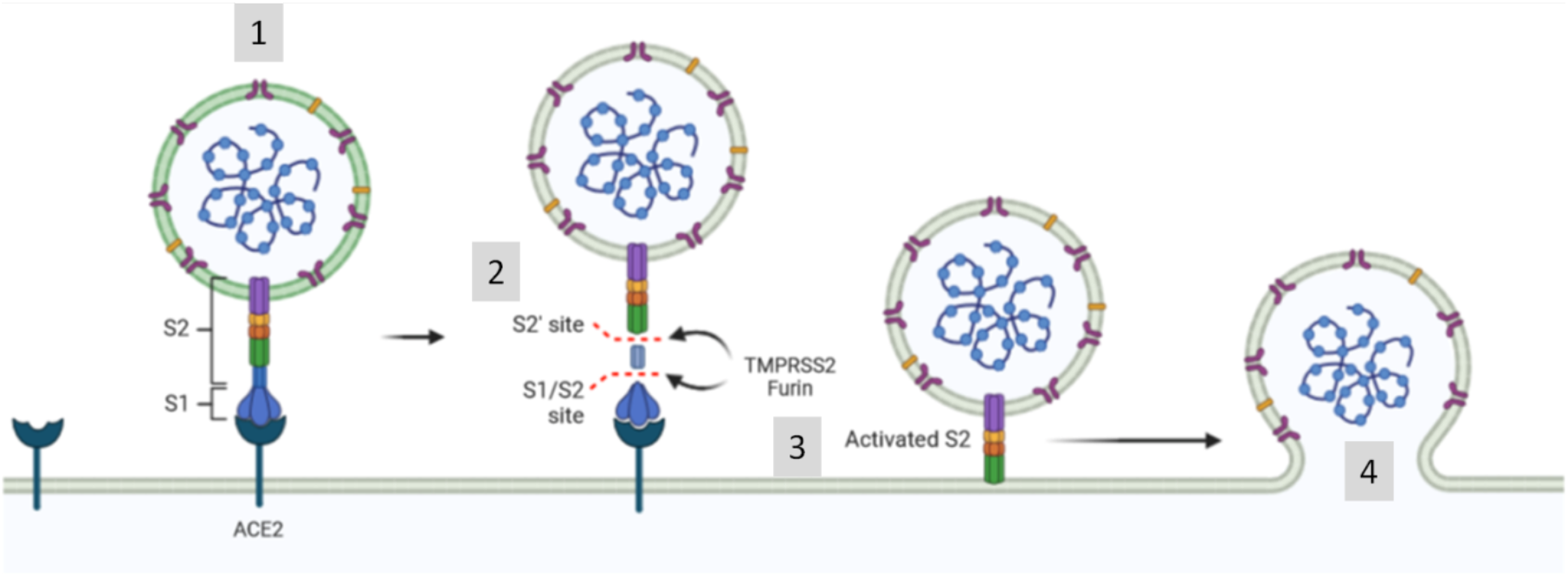
SARS-CoV-2 exploits the angiotensin-converting enzyme 2 (ACE2) to enter target cells. After receptor binding (1), the virus S protein is cleaved by proteases such as furin/TMPRSS2 into S1 and S2 subunits (2) that mediates S2-assisted fusion (3) and the release of the viral genome (4).

Viruses continually change through mutations, a mechanism responsible for the emergence of new variants. Particularly, RNA viruses have been reported to have elevated mutation levels as compared to DNA viruses [8,9]. In the surface protein of various viruses such as Ebola virus [10], Chikungunya virus [11] and the highly pathogenic avian influenza H5N1 [12], amino acid changes were found to significantly alter viral functions, often resulting in an increased transmissibility and/or mortality.

In the gene encoding S protein of SARS-CoV-2 various mutations were also reported [13,14] and recently, the United Kingdom (UK) and South Africa have faced a rapid increase in COVID-19 mediated by new variants that are named in the literature (VOC-202012/01 or VUI-202012/01 or B.1.1.7 for UK and 501Y.V2 or 20C/501Y.V2B.1.351 for South Africa). These variants harbor one or several non-synonymous spike mutations including amino-acid replacements at key sites in the spike RBD domain (K417N, E484K, N501Y for the African strain [15] and only N501Y in this region of the protein for the UK strain [16]). The proportion of these variants has increased rapidly and recent observations suggests that they are significantly more transmissible than previously circulating variants. However, it is still not fully known if the pathogenicity is increased, although some elements have been recently released for the UK strain with likely enhanced disease severity [17]. At present, it is still unclear why some individuals are more susceptible to virus infection and what could be the role of mutations on ACE2 or on the Spike protein. We attempt here to gain insight into this protein-protein interaction using various in silico approaches. We specifically focus on amino acid changes found in the South African and UK strains located in the Spike RBD domain with a special emphasis on the potential impact of the only three residues (K417, E484 and N501) that are located directly at the interface with ACE2.

## 2. Materials and Methods

We used the crystal structure of the Spike-RBD-ACE2 complex reported by Lan et al. [18] to map amino acid changes of the African and UK strains in 3D. We took into account the results of deep experimental mutational scanning of the spike RBD [19] to select the in silico tools that will be used to compute the energetics of the interaction and during the interactive structural analysis. COCOMAPS was used to analyze contacts at the biomolecular interface [20]. Flexibility of the ACE2-Spike complex was investigated with the CABS-flex package [21,22]. This approach uses a coarse-grained protein model to investigate flexibility so as to speed up the computations as compared to classical all-atom molecular dynamics and was shown to provide a similar overview of the system. Restraints were applied to take into account disulfide bonds present in the structure.

The global interaction energy differences between the initial Spike-ACE2 structure, the UK and South African strains were investigated with the SPServer [23]. This analysis uses split-statistical potentials to for instance investigate the interaction energy differences between native and mutant structures. Split-statistical potentials are knowledge-based potentials that consider the frequency of pairs of residues in contact, the nature of the amino acids and can include information about structural environment. In our computations, a Cbeta-Cbeta distance < 12 Å was used to instigate residues in contact. Different types of scores are reported. For instance, the score named PAIR is obtained by summing the potential of mean force with the corresponding subindex of each pair of interacting residues a, b, between the Spike and ACE2 proteins. It considers amino acid frequencies along distances. The pyDockEneRes software uses a different scoring function, it was used to provides interaction energy values of the protein-protein complex partitioned at the residue level [24].

The interactive structural analysis was performed with PyMol (Schrödinger product) and UCSF ChimeraX (32881101). Amino acid substitutions in the Spike protein were done using PyMol while a short energy minimization of each modified 3D complex was carried out with UCSF Chimera.

## 3. Results

The crystal structure of the Spike-RBD-ACE2 complex was analyzed using COCOMAPS and the results of the analysis indicates that 52 residues are involved in the interaction from the ACE2 side while 46 residues (see below) in the Spike RBD could play a role at the interface. About 1688 Å^2^ is buried upon complex formation and the buried polar surface represents about 58% of the interface while the non-polar buried surface involves about 42% of the interface. Thus, a variety of non-covalent interactions are found between the amino acids of both, the Spike protein and the ACE2 receptor. The identified residues on the Spike protein that have some contacts with ACE2 within a maximum distance of 8 Å from the ACE2 interface involve the following 46 amino-acids: R403, D405, E406, K417, Y421, N439, K444, V445, G446, G447, N448, Y449, Y453, R454, L455, F456, R457, Y473, Q474, A475, G476, S477, T478, E484, G485, F486, N487, C488, Y489, F490, P491, L492, Q493, S494, Y495, G496, F497, Q498, P499, T500, N501, G502, V503, G504, Y505, Q506 (here the strength or energetics of the interaction is not considered).

Protein flexibility was then investigated with CABS-Flex. The complex was simulated using the crystal structure as starting point **(Figure 3)**.

**Figure-3.**
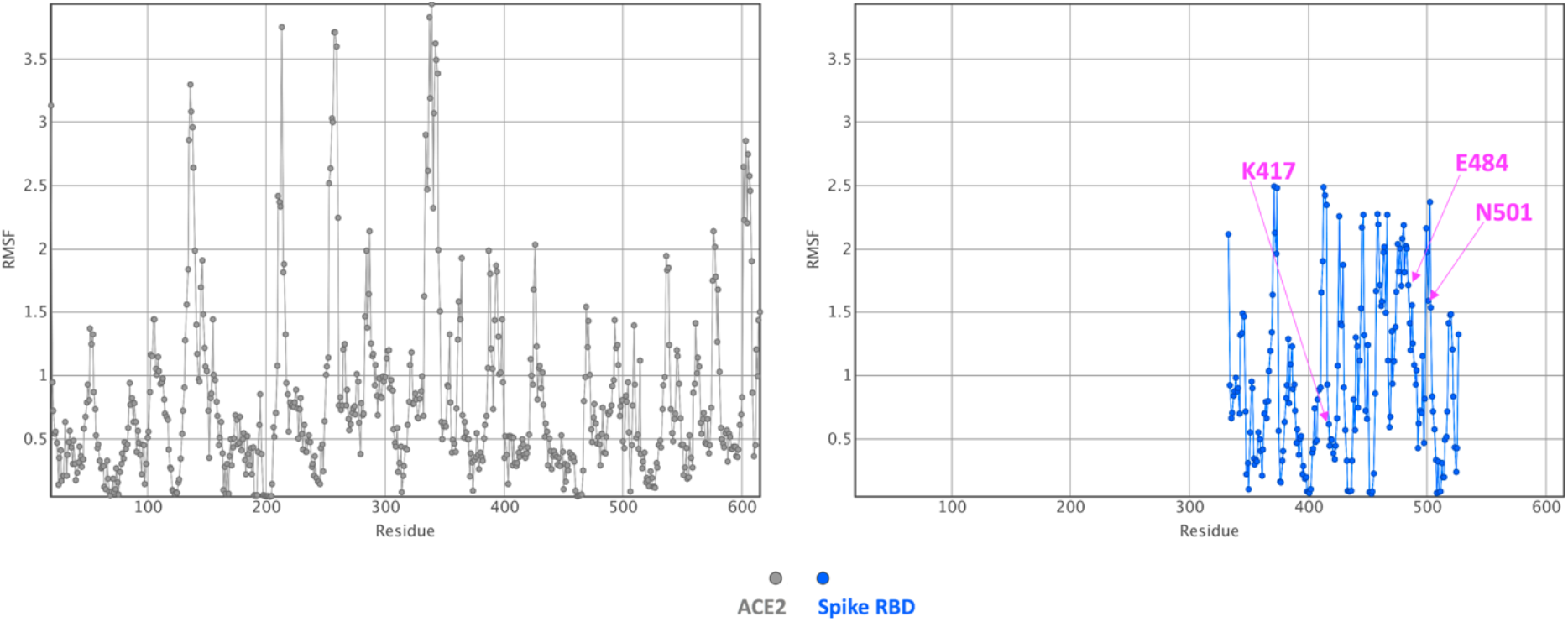
Fluctuation plot. The residue fluctuation profile (RMSF in Å) as computed with CABS-Flex for the ACE2-Spike RBD protein complex is shown (ACE2, in grey and the Spike domain, in blue). Substituted residues in this region of Spike in the UK strain (N501) and in the South African strain (K417N, E484K, N501Y) are shown in magenta.

The main global information that we obtained here is that many segments at the interface are relatively rigid, as expected, at least when the two proteins are bound. Yet, a peptide at the interface between Spike residues 475 and 487 is predicted to be more flexible (essentially a loop region) than the other interface regions. Some other segments on both proteins located relatively far away from the interface (essentially loops) are also predicted to be flexible. Simulations of the variant models do provide similar outputs (i.e., the substitutions did not enhance or reduce significantly the flexibility or rigidity of the interface, data not shown).

We then investigated the global binding scores for the original Spike-ACE2 structure as present in the PDB file and for the UK and South African strains with the SPServer. The PAIR score gives some insights about the energetics of the interaction. It was found to be more favorable for the UK strain (N501Y) than for the initial structure (N at position 501). But the predicted global interaction score between the Spike and ACE2 proteins seems less favorable for the South African strain (K417N, E484K, N501Y) than for the original input structure or for the UK strain.

In a following step, we predicted the interaction energy of the Spike RBD-ACE2 protein-protein complex partitioned at the residue level using pyDockEneRes. On the Spike RBD domain, the top residues that are predicted to have favorable interaction energy values with the region of ACE2 that binds the RBD are (from the most favorable according to this method, values of about −12 kcal/mol to less favorable but still with some contributions around −1 kcal/mol): F486, F456, Y505, Y489, R403, K417, Y453, R408, K444, N501, Q498, Q506, and A475 **(Figure 4)**. Some other residues are present at the interface on the Spike protein but the computed interaction energy values were outside our selected range (e.g., Spike Q493, Y449, L455, G485, or G496). Of interest, in the list of residues predicted to be important for the interaction with ACE2, Spike N501 (present in both the UK and South African strains) and Spike K417 (South African strain) are identified while E484 (the third substitution in this region of the Spike protein in the South African strain) is not predicted to be a major residue for the interaction. On ACE2, the top key predicted residues involved (from very favorable energy values around −7.7 kcal/mol to less favorable, around −1 kcal/mol) are: T27, Y83, K353, M82, D30, H34, D38, D355, E329, L45, and L79 (see Fig. 4). Here also, a few extra residues are visible at the interface (like ACE2 Y41 or K31) but the computed energy values were too weak considering the selected threshold value of about −1 kcal/mol.

**Figure 4.**
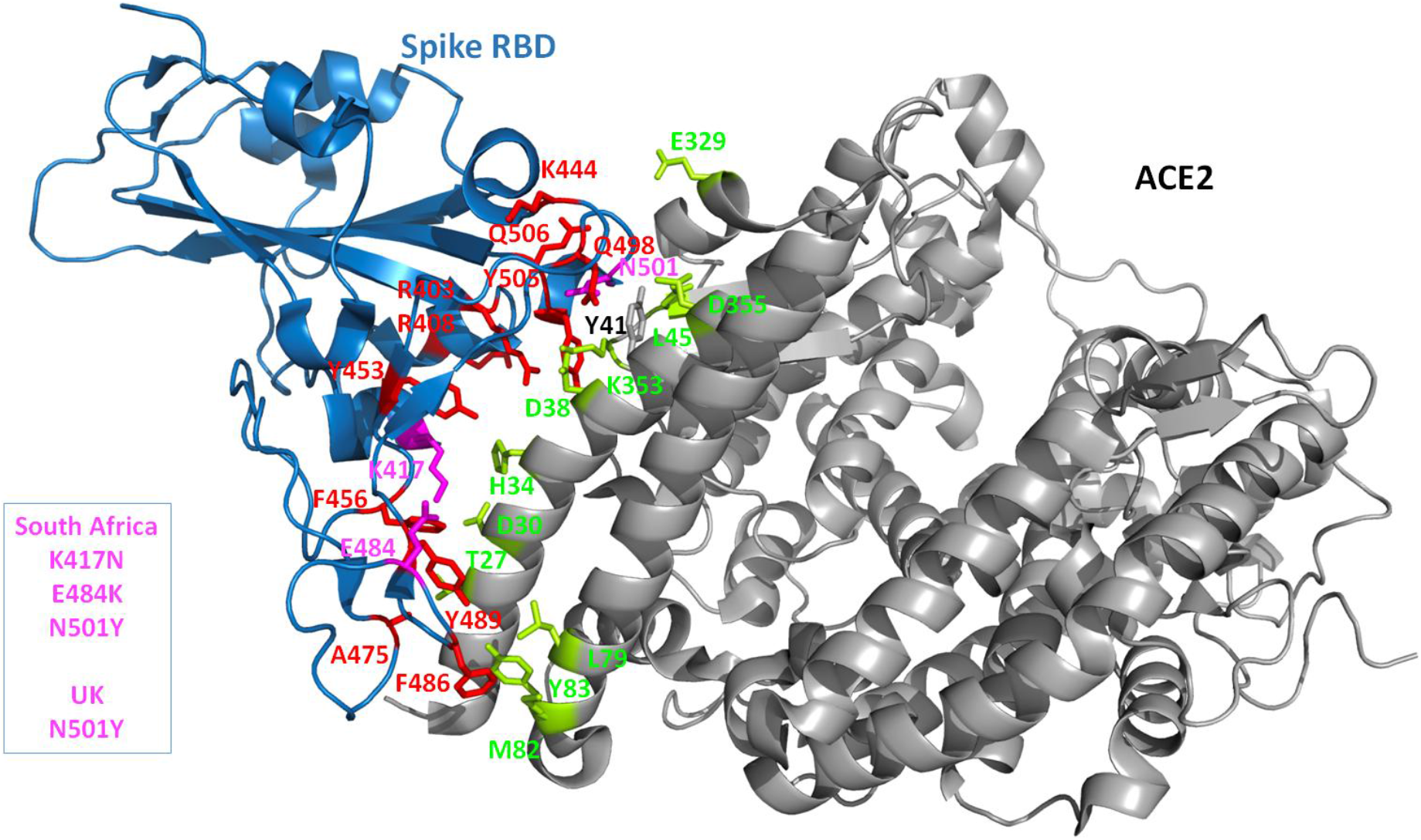
3D structure of the complex. The crystal structure of the Spike RBD and part of the ACE2 receptor interacting with the Spike is shown as cartoon diagram. The Spike domain is in blue while the ACE2 domain is in grey. The Spike side chains of residues predicted to be important (with our in silico protocol, see the Method section) for the interaction with ACE2 are shown in red. The UK and/or South African strains are highlighted by coloring the amino acid changes in this region in magenta. All the Spike side chains shown should have favorable interaction energies with ACE2 but E484 that is predicted to have very weak or unfavorable interactions). On the ACE2 side, the side chains that are predicted to contribute favorably to the interaction with the Spike RBD are shown in lemon. Only ACE Y41 seems to have very limited interactions with the Spike protein while, when the Spike protein carries a Y at position 501, ACE2 Y41 has then some favorable interaction energy values with the Spike.

If we then consider the three residue replacements of interest in the present analysis (N501Y, K417N, E484K), we observe that N501 (UK and South African strains) is solvent exposed on the free Spike protein and becomes essentially buried upon ACE2 binding. N501 is located in a loop structure and could be replaced by a Y without creating folding problems. Some flexibility is present there suggesting that a larger Y residue could be accommodated (**Figure 3**). Some weak hydrogen bonds are possible with ACE2 K353 (a residue found to have some contributions with the pyDockEneRes tool) and ACE2 Y41 (not listed in our analysis as the computed contribution with this method is too weak considering our threshold). The contribution of N501 to the interaction with ACE2 is predicted to be around −1.6 kcal/mol but when a Y is present at this position, the variant residue has a predicted interaction energy with ACE2 of about −3.5 kcal/mol. On the graphics display, we note that the newly introduced Y501 side chain should make many favorable interactions with ACE2, like for example with ACE2 Y41 via a Pi-stacking, with ACE2 K353 via a cation-Pi interaction and possibly with ACE2 D38 via a hydrogen bond. This type of non-covalent interactions (Pi-stacking and cation-Pi) in the variant should be very favorable for the interaction and much stronger in strengths as compared to the initial N residue. It is important to note that in fact such energetics terms are in general underestimated or not considered by many scoring functions as very difficult to calibrate.

Spike K417 is located at the N-term of a short helix, it is solvent exposed and becomes partially buried upon binding to ACE2. It is in a somewhat rigid region of the interface (**Figures 3 and 4**). It can however be replaced by N (South African strain) without creating folding problems and should not destabilize the Spike protein. It forms a salt-bridge with ACE2 D30 and should contribute favorably to the interaction as its predicted energy value is around −3 kcal/mol. Upon replacement by a N, the contribution of position 417 to the interaction with ACE2 is less favorable (loss of salt-bridge) than with the K and is computed to be −1.3 kcal/mol. Some weak hydrogen bonds could still be possible, most likely with ACE2 H34 and D30.

E484 is located in a loop structure and is solvent exposed. This segment is somewhat more flexible than the remaining regions of the interface (**Figures 3 and 4**). It remains solvent exposed upon binding to the ACE2 protein. Its contribution to the interaction energy is predicted to be relatively weak in spite the presence of the nearby ACE2 K31 (most likely because of the distance between the two charge centers is around 4.4 Å (slightly higher than 4 Å or less, commonly found in energetically stronger salt-bridges) and as the interaction is fully solvent exposed). E484 could be replaced by a K in the Spike protein without creating folding problem. Yet, a K at this position in the ACE2-Spike complex is predicted to be less favorable than a E with regard to the interaction with ACE2 (about +1.9 kcal/mol, possibly the proximity with ACE2 K31).

## 4. Discussion

Amino acid changes in a protein can have numerous impacts, from folding problems to modulation of intermolecular interactions with ligands or protein partners. Several key structural properties are known to play a role here, such as the type of amino acid substitution (conservative replacement or not), the location in the protein structure (e.g., the change takes place in a loop or in secondary structure elements, the residue is buried or solvent exposed), the residue is inside a catalytic site or within a protein-ligand interface, the replaced residue is in a flexible or rigid region among many others [25–32]. Different types of computational tools (https://www.vls3d.com/), all with strengths and weaknesses, can be used to investigate the possible impacts of amino acid replacements on the structure and function of a protein when a 3D structure is available or can be predicted [25–32]. It is recommended to use different tools that apply different types of algorithms so as to gain some “consensus” insights about the amino-acid replacement [25]. Further, when experimental data are available, it is obviously of interest to select methods that can at least reproduce such data. The present investigation benefits from the recently published Spike mutagenesis study and the impacts on ACE2 binding [19]. Among the experimental data reported in that study, we were particularly interested in the measured affinity of the Spike N501F protein mutant with ACE2 as it involves residue 501 and as the affinity was assessed in a purified binding assay (most other affinity evaluation were performed using high-throughput affinity measurements that could be less accurate than measurements carried out in purified systems). Experimentally, it was found that the Spike N501F protein has increased affinity for ACE2. Interestingly, most in silico tools that we tested to compute stability changes at protein interfaces (ΔΔG) could not reproduce this measurement. We noted that the SPServer and pyDockEneRes tools were able to output data consistent with this high-quality N501F experimental result. Our selection does not mean that these in silico approaches will systematically performed better than the other methods in all cases but that they are interesting for this specific study. Further, while these tools have been assessed extensively by the authors, we still decided to compare the results of these two methods on five protein-protein interactions selected from the SKEMPI database [33] (thus with measured experimental stability changes upon mutations) to gain confidence in the meaning of the computed scores (data not shown). That is, all methods that predict such stability changes are compared with experimental values and correlations between the prediction and the measured values are reported. This is of importance but equally important is to gain insight about the type of substitutions that are well covered by the methods (e.g., a method recognizes or not a salt-bridge, the energetics make sense as compared to what is seen on the computer display, etc. [25].

Several crystal structures of the ACE2 and SARS-CoV-2 receptor-binding domain (RBD) have been reported (PDB IDs: 6LZG, 6M17, 6M0J) [34] to help the investigation the specific residues at the interface. For example, several polymorphisms in the ACE2 gene have been proposed to reduce the affinity toward the Spike protein, with subsequent lower susceptibility to infection [35]. Also, structural and in silico analyses of ACE2 polymorphisms have been carried out on the ACE2 region that directly interacts with the Spike glycoprotein [36]. In that study, the authors found, for example, that two substitutions in ACE2 at positions 19 and 26 could modulate the affinity for the Spike protein (ACE2 position 19, fully solvent exposed, the S19P is common in African people and the substitution was suggested to protect individuals (reduced affinity for Spike) and a cleavage site of the ACE2 precursor; ACE2 position 26, fully solvent exposed, the K26R is common in European people, the mutation may increase affinity of ACE2 towards Spike). However, in our hands, it would seem that the ACE2 K26R may only have minor roles with regard to Spike binding. In our investigation, residue K26 was not found to be a major player that strongly contributes to the stability of the interface because it is at about 8 Å from the Spike protein in all the available experimental structures of the complex and in fact tends to point away from the Spike surface (of course some side chain flexibility is likely but in the available X-ray structures, around ACE2 K26 we note the Spike N487 side chain, at about 7.8 Å, much too far for hydrogen bonds and the Spike K417 side chain, at about 7Å with possibly some electrostatic repulsion). Some long distance electrostatic interactions may play a role in the pre-orientation of the two binding partners when ACE2 K26 is replaced by arginine (e.g., could we have some improved hydrogen bond networks due to the amino-acid substitution with water molecules at the interface?). Another alternative could be that, as ACE2 K26 makes a salt-bridge with ACE2 E22 and polar interactions with ACE2 N90, the K26R substitution stabilizes locally this region of ACE2 (with possibly creating another salt-bridge with the nearby ACE2 E23 not possible when ACE2 at position 26 is a K, data not shown) and contributes to a more favorable change in free energy of binding with the Spike protein.

Some additional investigations are needed to clarify these observations. Some other studies investigating genetic variations in ACE2 suggest that amino acid changes in regions that are relatively far away from the interaction site for the Spike protein could also alter the recognition via various molecular mechanisms [37].

The main focus of the present study is the Spike protein. For this macromolecule, some mutations have also been reported; some could increase the infectivity of the virus. For example, the Spike D614G substitution [14,38] seems to increase affinity although this residue is not present at the Spike-ACE2 interface but at a Spike protomer-protomer interface. Here, we were interested in the analysis of one or three amino acid substitutions as found in the UK strain and South African strain, respectively, with a special emphasis on residues that are directly present on the Spike receptor-binding domain, at the interface with ACE2. We wanted to gain some insights about the putative impacts of these substitutions on the interaction with the ACE2 receptor. Our in silico results were also investigated in the light of recently reported experimental mutagenesis data [19]. Experimentally, amino acid can be substituted and the expression levels monitored (e.g., notion of stability problem if for instance the mutant protein is expressed at very low level). Such behavior can be estimated in silico. Further, when a mutant protein is expressed, its interaction with a protein partner can be evaluated. This event can also be investigated to some extent in silico. In all cases, it is important to note that both, experimental data and in silico predictions can be misleading and as such results should be investigated with cautions.

In the previously reported mutagenesis study mentioned above [19], it was found that the replacement of Spike N501 by a F enhances ACE2 affinity while having no effects on expression. This observation is expected for the N501Y substitution although the measurements were not performed in purified system. Overall, these experimental data are in agreement with the present computer analysis. The replacement of K417 by N (South African strain) seems relatively neutral in term of protein expression level while not very favorable for the interaction with ACE2 [19], also in agreement with our computer evaluation. The E484K replacement was not found to change the expression level, suggesting, as observed in our structural analysis, that a K could be present in this position of the Spike protein. Yet, the substitution with a K seems experimentally to slightly enhance the affinity towards the ACE2 receptor while we computed that a K at this position should be less favorable than a E. The differences between the computed values and the experimental observation could be due to inaccuracies in the scoring functions and unexpected flexibility at the interface (yet, introducing flexibility with in silico approaches can lead to nonsense energy values). It is also important to note that the substitutions in the South African and UK strains outside the interface area with ACE2 could play a role in affinity and/or on transmission/infectivity. Yet, when focusing only at the Spike RBD-ACE2 interface, the present in silico predictions and interactive structural analysis suggest that the key player residue in term of enhanced affinity is the N501Y replacement. Our observation applies essentially to the UK strain while additional work is required to gain insights over the South African strain because in our hand, looking only at the interface, it would seem that the K417N and E484K substitutions are not favorable for the interaction with ACE2. For the time being, we do not know if the N501Y alone could counterbalance the predicted unfavorable Spike K417N and E484K amino acid replacements. Considering the clinical observations about the South African strain and potentials greater risks relative to the UK strain, as it is definitively unclear that the affinity between the Spike and ACE2 is significantly enhanced (according to our predictions), then, danger is due to some amino acid changes (e.g., Spike residue 484 in this region of the protein and possibly elsewhere) that impede recognition by neutralizing antibodies [39,40]. For the UK strain, could it be that the enhanced affinity to ACE2 not only increases transmissibility but also disease severity (not only due the increased number of infected individuals) as suggested after comparisons between SARS-CoV-2 and other human coronaviruses [41]?

## 5. Acknowledgements

AMK and BV have been supported by Inserm.

## 6. Credit Author Statement

**Bruno Villoutreix:** Methodology, Analysis, Investigation, Writing and Editing. **Vincent Calvez**: Editing. **Anne-Genevieve Marcelin: Editing**. **Abdel-Majid Khatib:** Writing and Editing.

